# The methyl cycle is a conserved regulator of biological clocks

**DOI:** 10.1101/653667

**Authors:** Jean-Michel Fustin, Shiqi Ye, Christin Rakers, Marijke Versteven, Samantha J. Cargill, T. Katherine Tamai, Yao Xu, Maria Luísa Jabbur, Rika Kojima, Melisa L. Lamberti, Kumiko Yoshioka-Kobayashi, David Whitmore, Ryoichiro Kageyama, Takuya Matsuo, Ralf Stanewsky, Diego A. Golombek, Carl Hirschie Johnson, Gerben van Ooijen, Hitoshi Okamura

## Abstract

The methyl cycle is a universally conserved metabolic pathway operating in prokaryotes and eukaryotes. In this pathway, the amino acid methionine is used to synthesize S-adenosylmethionine, the methyl donor co-substrate in the methylation of nucleic acids, histone and non-histone proteins and many other molecules within the cell. The methylation of nucleic acids and proteins is the foundation of epigenetic and epitranscriptomic regulations of gene expression, but whether the methyl cycle centrally regulates gene expression and function by controlling the availability of methyl moieties is poorly understood.

From cyanobacteria to humans, a circadian clock that involves an exquisitely regulated transcription-translation-feedback loop driving oscillations in gene expression and orchestrating physiology and behavior has been described. We reported previously that inhibition of the methyl cycle in mammalian cells caused the lengthening of the period of these oscillations, suggesting the methyl cycle may indeed act as a central regulator of gene expression, at least in mammals. Here, we investigated whether the methyl cycle, given its universal presence among living beings, regulates the circadian clock in species across the phylogenetic tree of life.

We reveal a remarkable evolutionary conservation of the link between the methyl cycle and the circadian clock. Moreover, we show that the methyl cycle also regulates the somite segmentation clock, another transcription-translation negative feedback loop-based timing mechanism that orchestrate embryonic development in vertebrates, highlighting the methyl cycle as a master regulator of biological clocks.

**SIGNIFICANCE STATEMENT:** Here we reveal that the methyl cycle, a universal metabolic pathway leading to the synthesis of S-adenosylmethionine, the methyl donor co-substrate in virtually all transmethylation reactions within the cell, is a conserved regulator of biological clocks. These discoveries highlight the methyl cycle as a metabolic hub that regulates gene expression via the availability of methyl moieties for the methylation of nucleic acids, proteins and many other molecules with the cell.

## INTRODUCTION

Methylation reactions start with the metabolization of methionine into *S*-adenosylmethionine, or SAM: the universal methyl donor co-substrate in the transmethylation of nucleic acids, proteins, carbohydrates, phospholipids and small molecules. During the methylation process, SAM is converted into adenosylhomocysteine (SAH) that is rapidly hydrolysed into homocysteine to prevent competitive inhibition of methyltransferase enzymes by SAH due to its close structural relatedness to SAM. Homocysteine is detrimental to physiology (*1*), and is recycled back into methionine or used for glutathione synthesis via cysteine and cystathionine in the trans-sulfuration pathway. The ratio SAM/SAH is known as the methylation potential: a measure of the tendency to methylate biomolecules (*2–4*). In plants, fungi, prokaryotes and other microorganisms, the methyl cycle is used both as a source of SAM, as well as a source of *de novo* methionine synthesis for growth. In a wide variety of eukaryotes, however, methionine is an essential amino acid that must be obtained from food, especially in periods of growth, the methyl cycle in these organisms ensuring only the flow of one-carbon units to SAM.

Three enzymatic activities are required to keep the methyl cycle running. 5-methyl-tetrahydrofolate or betaine are used as a source of methyl moieties to regenerate methionine (Met) from homocysteine (Hcy). The enzyme *Methyltetrahydrofolate-homocysteine S-methyltransferase* is universal but *Betaine-homocysteine methyltransferase* is present only in some vertebrate tissues, mainly in the liver. Likewise, *Adenosylmethionine synthetase* (SAM synthesis from Met) and *adenosylhomocysteinase* (Hcy synthesis from SAH) are virtually universal. There is considerable divergence in methyltransferases (MTases), and the phylogeny of five structural classes of MTases is unclear. The oldest MTases represents the RNA methyltransferases, appearing in the early forms of life evolving on Earth in an RNA World (*5*). Methyl cycle metabolites are thought to have been present in the prebiotic world, since methionine can be created by a spark discharge and is proposed to be an intermediate in the prebiotic synthesis of homocysteine, which was also found among organic molecules synthesized in the 1972 Miller experiment (*6*). Prebiotic chemistry might have involved methylation by reaction with formaldehyde, an abundant prebiotic organic molecule, to prime the evolution of biological methylation reactions (*7*).

Similarly, an endogenous circadian clock evolved to anticipate the daily cycles of light and darkness has been found in many organisms, from cyanobacteria to humans. Transcription-translation feedback loops (TTFLs) of “clock genes” directly or indirectly regulating their own transcription underlie many functions of the clock, and drive oscillations of output genes controlling physiology and behavior. Some molecular components of the clock are remarkably conserved in metazoan, notably the genes *Clock* and *Period*, coding for transcription factors, with *Clock* activating the transcription of *Per* and *Per* inhibiting its own transcription.

In 2013, we reported that disruption of the methyl cycle by inhibitors of *Adenosylhomocysteinase* (AHCY), causing an accumulation of SAH known to lead to a general inhibition of transmethylations, strongly affected the circadian clock in mouse and human cells (*8*). We now show that the link between the methyl cycle and the circadian clock we first uncovered in mammals has been conserved during more than 2.5 billion years of evolution. We further reveal a similar link between the methyl cycle and another TTFL called the somite segmentation clock that underlies the development of early body plan in vertebrates, indicating that the methyl cycle is a key regulator of biological clocks.

## RESULTS

### AHCY is a remarkably conserved key enzyme in the methyl cycle

The use of carbocyclic adenosine analogues such as Deazaneplanocin A (DZnep) as inhibitors of AHCY was established more than 30 years ago (*9–11*). The reaction catalyzed by AHCY is the cleavage of SAH to adenosine and L-homocysteine, DZnep inhibiting this reaction by occupying the adenosine binding site. The crystal structure of human (14), mouse (15) and yellow lupin (*Lupinus luteus*) (16) AHCY complexed with adenosine or analogues have been described, and insights into its catalytic activity have been obtained (*17–20*). AHCY has been reported to be one of the most evolutionarily conserved proteins (21), but experimentally-determined structures of AHCY with DZnep remain to be described for most organisms investigated here. A full-length multiple sequence alignment (Fig. S1) and homology modelling of AHCY from human to cyanobacteria (Fig. 1a and Movie S1) revealed high sequence and predicted tertiary structure conservation. Amino acids contributing to the DZnep binding site showed at least 88% identity between all eukaryotic AHCY sequences, and 78% between human and bacterial sequences, resulting in the DZnep binding site to be virtually identical in all organisms investigated (Fig. 1b, c and Fig. S2). Moreover, amino acids that were reported as crucial for the activity of rat AHCY (His55, Asp130, Glu155, Lys186, Asp190, and Asn191 (*20*)) are perfectly conserved (Fig. S1). In line with the above, molecular docking simulations of AHCY with adenosine or DZnep revealed comparable ligand binding conformations and estimated binding free energies for all organisms (Fig. 1d and S2). Together these observations strongly support the use of DZnep as a valid approach to test the effects of methyl cycle inhibition on clocks across phyla.

**Fig. 1:**
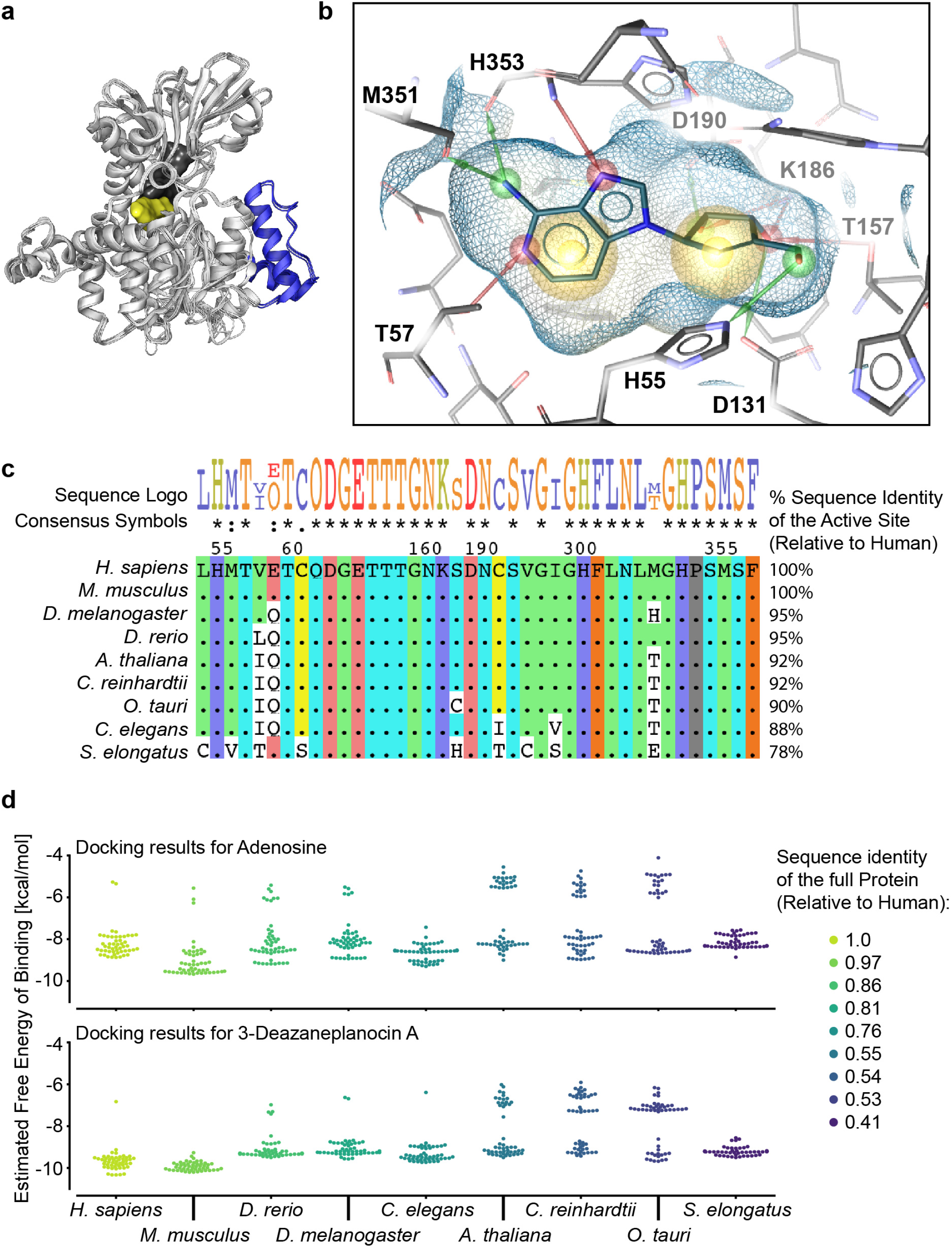
Adenosylhomocysteinase is a highly conserved protein. **(a)** Structural superposition of AHCY from the 9 organisms investigated here, using human (1LI4), mouse (5AXA) or lupin (3OND) crystal structures as templates. The blue loop is specific to plants and green algae; DZnep is shown in yellow, NAD^+^ in grey. See also Movie S1. **(b)** Docking simulation of human AHCY with DZnep, based on the 1LI4 crystal structure of human AHCY complexed with Neplanocin A, an analogue of DZnep. The amino acids involved in DZnep binding are indicated with their position. See Fig. S2 for docking simulations of DZnep to AHCY from other organisms. Red and green arrows are hydrogen bonds, yellow spheres are hydrophobic effects. The estimated free energy of binding for depicted DZnep docking conformation was −9.87 kcal/mol. **(c)** Discontinuous alignment of amino acids contributing to the binding of DZnep, using the human sequence as a reference and with sequence identities shown on the right. When amino acids are identical to human, a dot is shown in the alignment. The sequence logo on top is a graphical representation of the conservation of amino acids, with the consensus symbols below (* = fully conserved residue, : = conservation of strongly similar properties,. = conservation of weakly similar properties). The positions of selected conserved amino acids are given for the human sequence. **(d)** Molecular docking simulations of AHCY with adenosine (top) or DZnep (below) showing comparable binding free energies in all organisms. Colors represent full sequence identities, relative to human.

### The circadian clock and methyl cycle are linked in vertebrates

We first tested the effects of DZnep on the mammalian clock, in a human osteosarcoma cell line stably transfected with a luciferase reporter vector for the core clock gene *Bmal1* (22) and in mouse embryonic fibroblasts from PER2::LUC knock-in mice expressing a fusion between the endogenous core clock protein PER2 and LUCIFERASE (23). These cells are the gold standard for measuring circadian parameters *in vitro* since they allow the oscillating expression of the genes *Bmal1* or *Per2* to be followed in real-time. In human (Fig. 2a) and mouse (Fig. 2b) cells, DZnep dose-dependently increased the period, up to a maximum of ∼40 hours and ∼30 hours, respectively, with 100 μM DZnep.

**Fig. 2:**
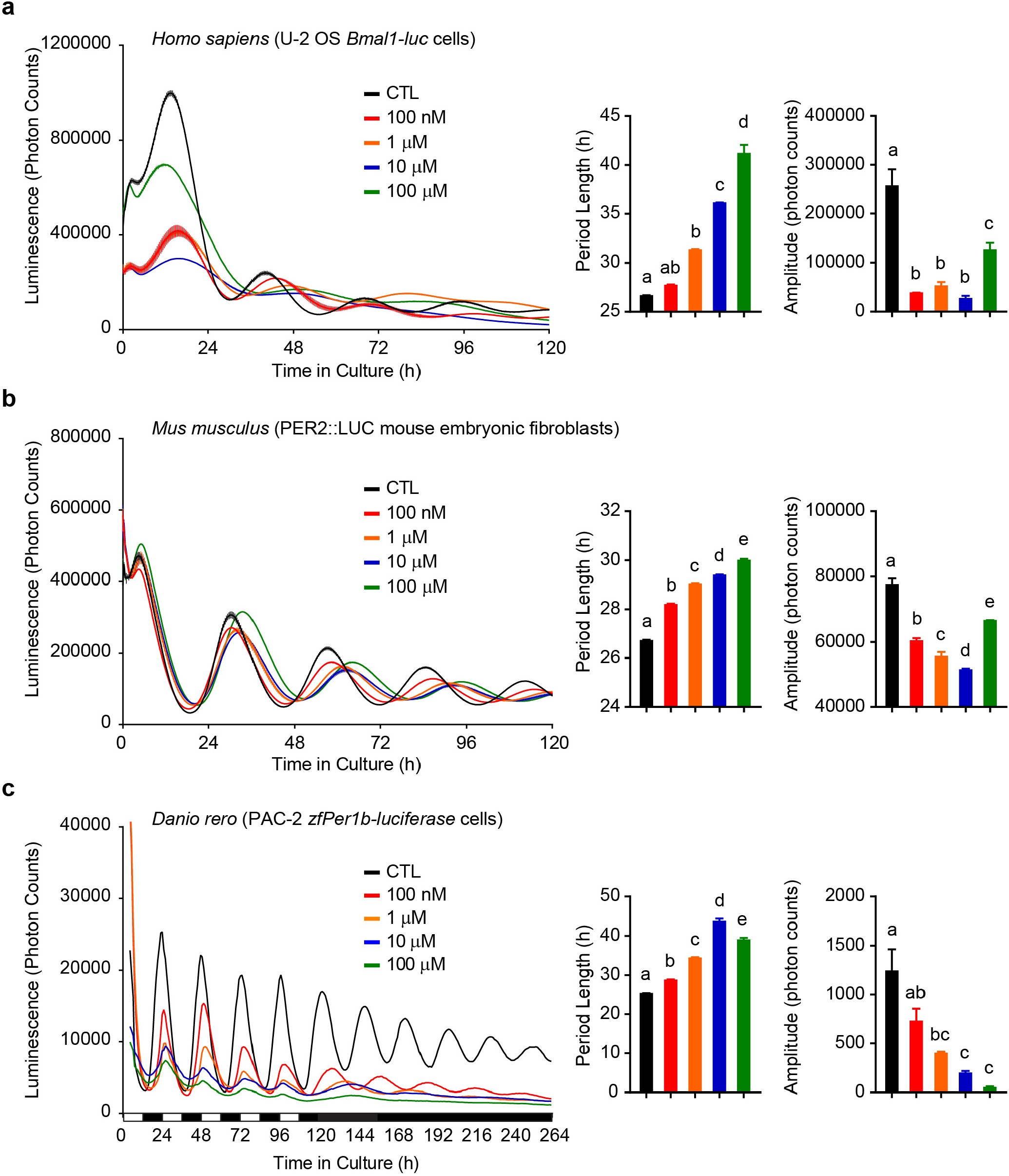
The link between the methyl cycle and the clock is conserved in vertebrates. **(a)** Left panel shows mean luminescence +/- SEM of human U-2 OS cells stably transfected by a *Bmal1*-luc reporter and treated with increasing concentrations of DZnep. Middle panel shows mean +/- SEM of period, n = 3. Right panel shows mean +/- SEM of amplitude, n = 3 dishes. Same analyses were performed with PER2::LUC mouse embryonic fibroblasts, n = 3 dishes **(b)**, and PAC- 2 zebrafish cells stably transfected with a *Per1b*-luciferase reporter, n = 4 dishes, mean +/- SEM **(c)**. These experiments were independently reproduced at least three times. All bar graphs analyzed by One-Way ANOVA followed by Bonferroni post-hoc test; all indicated comparisons at least *p* < 0.05.

We next tested DZnep on a non-mammalian cell type commonly used for circadian studies, the zebrafish embryonic PAC2 cell line and revealed a potent effect of the drug both on circadian period and amplitude (Fig. 2c), as in mammals. Since the circadian clock in these cells is directly light-sensitive and can be entrained by light-dark cycles, we also tested whether the entrainment of the cells to light was affected by DZnep before release into constant darkness (DD). Indeed, a strong effect on entrainment was observed (Fig. 2c). The gene *Per1b* normally peaks at dawn but was severely blunted and delayed in DZNep-treated cells, even at the lowest concentration of DZnep. In DD, period lengthened dramatically to a peak value of over 40 hours at 10 μM, that was slightly lower at 100 μM. To conclude, period and amplitude of circadian oscillations of clock gene expression are strongly affected by DZnep in all three vertebrate cell systems.

### The circadian clock and methyl cycle are linked in invertebrates

As models for invertebrates, we selected the fruit fly *Drosophila melanogaster* and the nematode *Caenorhabditis elegans*. Flies have been instrumental in molecular circadian biology, allowing the 2017 Nobel laureates and their co-workers to establish the canonical TTFL model underlying circadian oscillations in metazoan (24–26).

In contrast, *C. elegans* is a relatively new model in circadian biology, the general principles governing its circadian clock remaining largely unidentified (*27, 28*). Circadian oscillations in the expression of the *sur-5* gene were described a few years ago and used for the real-time monitoring of the nematode’s circadian molecular rhythms by luciferase reporter (29).

As observed in vertebrates, DZnep caused dose-dependent period lengthening in *D. melanogaster* haltere cultures from transgenic luciferase reporter TIM-LUC flies (Fig. 3a), and a significant effect of 100 μM DZnep was also observed in freely moving nematodes. Period changes in flies were of lesser magnitude than observed in vertebrates. Nevertheless, these results indicate that the link between the circadian clock and the methyl cycle is conserved in invertebrates.

**Fig. 3:**
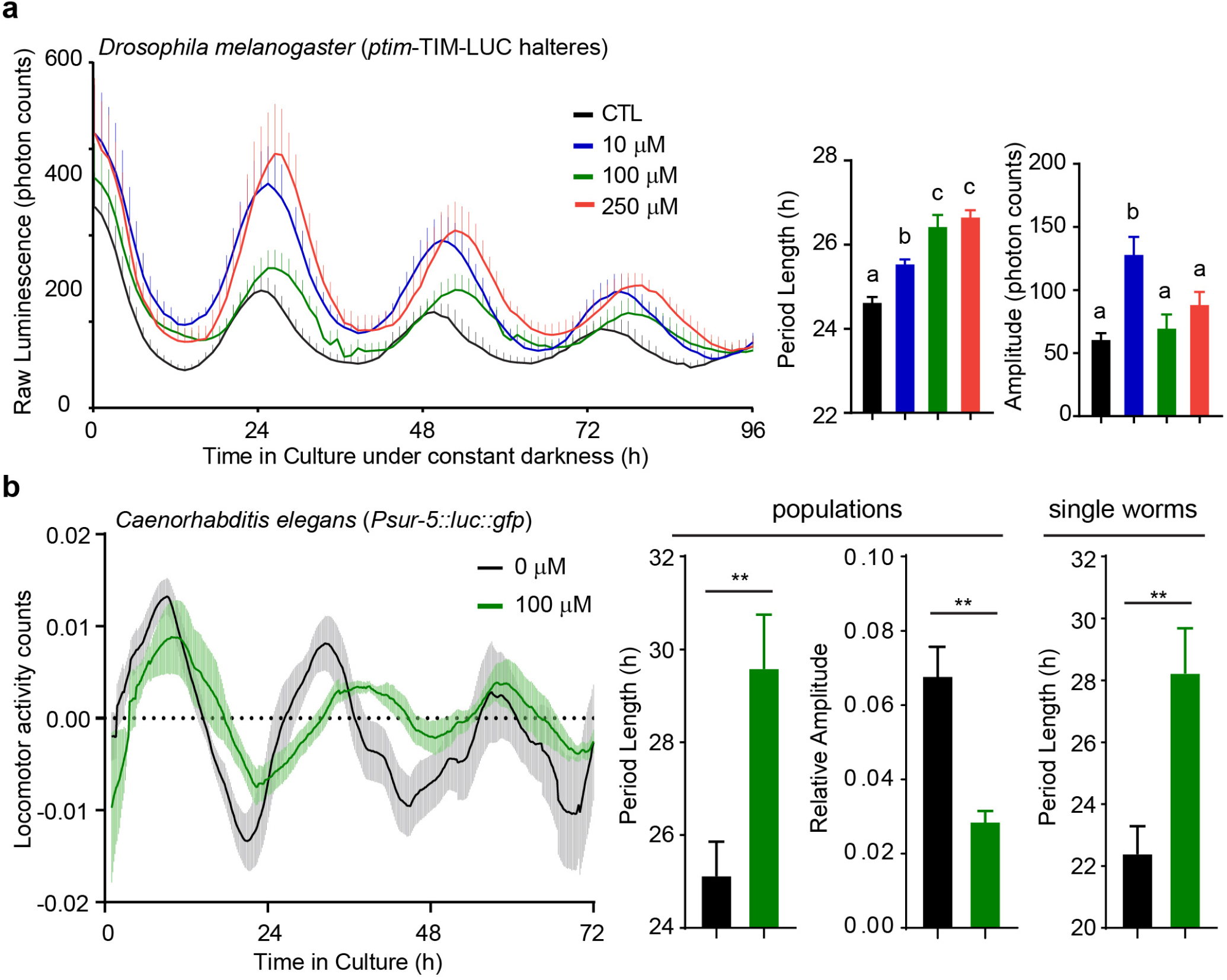
The link between the methyl cycle and the clock is conserved in invertebrates. **(a)** Real-time luminescence from halteres in culture dissected from *ptim*-TIM-LUC male flies. Luminescence traces show mean +/- SEM of n = 8 halteres, with only the upper segment of the error bars shown for clarity. Middle and right panel shows mean +/- SEM of period and amplitude, respectively, for n = at least 8 halteres for each treatment group. All bar graphs analyzed by One-Way ANOVA followed by Bonferroni post-hoc test; all indicated comparisons at least *p* < 0.05. **(b)** Locomotor activity counts from detrended luminescence measurements of freely moving *Caenorhabditis elegans* populations of ∼100 nematodes treated with vehicle or 100 μM DZnep, showing mean +/- SEM of n = 10 populations. In the middle, population mean +/- SEM of period (left) and amplitude (right) compared by Student *t*-test; **, *p* < 0.01; n = 10 populations treated with vehicle and n = 6 treated with DZnep. The right panel shows mean +/- SEM of period obtained from single nematodes in isolation, n = 21 controls and 22 for DZnep, analyzed by Student *t*-test, **, *p* < 0.01.

### The circadian clock and methyl cycle are linked in plants and algae

Fig. 1 and S1 show that a plant- and green algae-specific region exists in AHCY, from amino acids 151 to 191, which is involved in the interaction of AHCY with adenosine kinase and cap methyltransferase, and required for nuclear targeting of the enzyme (*30, 31*). Despite this insertion, however, the domains for SAH and NAD^+^ binding are remarkably conserved.

The most commonly used model organism to study circadian rhythms in land plants is *Arabidopsis thaliana*. We thus tested the effects of DZnep on luminescent rhythms reporting the expression of the plant evening gene TOC1 in protoplasts (Fig. 4a), and extended our investigations to aquatic unicellular green algae (Fig. 4b and c). *Ostreococcus tauri* and *Chlamydomonas reinhardtii* represent two different classes of unicellular green algae that have been successfully used in circadian studies and constitute great models to investigate cell-autonomous metabolic processes (*32–36*). We tested increasing concentration of DZnep on *Arabidopsis* and algal cell types and observed an increase in period length and a decrease in amplitude almost identical to the results obtained in vertebrates. In conclusion, the effects of DZnep treatment on transcriptional rhythms are also conserved in the plant kingdom. These results are especially significant given unicellular algae and humans are separated by more than 1 billion years of evolution.

**Fig. 4:**
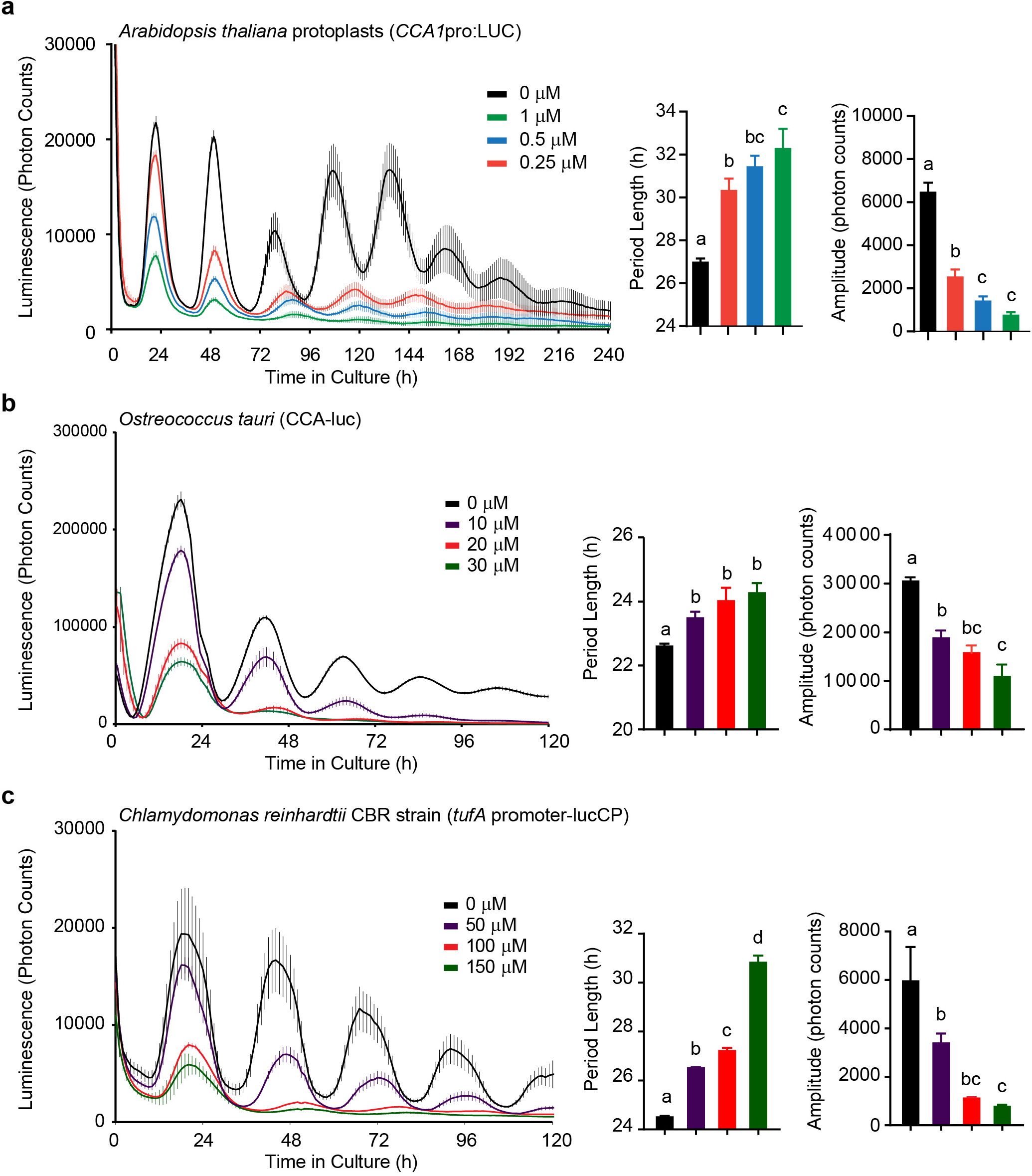
The link between the methyl cycle and the clock is conserved in plants. **(a)** Left panel shows mean luminescence +/- SEM of *Arabidopsis thaliana* protoplasts bearing a *CCA1*pro:LUC reporter construct, n = 8 wells per treatment, treated with different concentration of DZnep. For comparison between different runs, traces were aligned in relation to the first peak. Middle panel shows mean +/- SEM of amplitude, n = 8. Right panel shows mean +/- SEM of amplitude, n = 8. **(b)** Same as (a) but with *Ostreococcus tauri* cells carrying a CCA1-LUC reporter, n = 8 wells. Middle panel shows mean +/- SEM of period, n = 8 wells. No significance was observed between 10, 20 and 30 μM, but the significance compared to 0 μM became stronger, i.e. *p* < 0.05, *p* < 0.001, *p* < 0.0001, respectively, indicating dose-dependent effects. Right panel shows mean +/- SEM of amplitude, n = 8 wells. **(c)** Same as (b) but with *Chlamydomonas reinhardtii* CBR strain cultures stably-transfected with a *tufA* promoter-lucCP reporter, n = 5 wells. Middle panel shows mean +/- SEM of period, n = 5 wells. Right panel shows mean +/- SEM of amplitude, n =5 wells. All bar graphs analyzed by One-Way ANOVA followed by Bonferroni post-hoc test; all indicated comparisons at least *p* < 0.05.

### The circadian clock and methyl cycle are linked in prokaryotes

Eukaryotic circadian clocks involve a complex TTFL system that requires regulated gene expression at every level from transcriptional to post-translational steps: DNA methylation and the histone code, RNA processing, translation efficiency, phosphorylation and other protein modifications (*37*). In contrast, a biochemical oscillator in the cyanobacteria *Synechococcus elongatus* can independently generate a ∼24-hour rhythm. Three proteins, Kai-A, -B and -C, form a nanocomplex that regulates two key activities of KaiC: ATPase and autophosphorylation. The result is an autonomous and self-sustained phosphorylation-based oscillator ticking with a period close to 24-hours. In a cellular context, this non-transcriptional oscillator controls transcriptional outputs that in turn add robustness to the biochemical oscillator via TTFLs (*38–42*).

Like all organisms, cyanobacteria synthesize SAM, required for essential transmethylations. In cyanobacteria we observed a more complex response to DZnep than in other organisms, perhaps due to the presence in these cells of a self-sustained biochemical oscillator that drives the TTFL system. While circadian period lengthened in response to low concentrations of DZnep, the amplitude increased (Fig. 5a). These observations were confirmed using a different reporter strain of *S. elongatus* (Fig. 5b). Probing the methyl cycle-clock relationship further revealed that intermediate concentrations of the drug caused no significant effect on period but still significantly increased the amplitude (Fig. S3). At 200 μM however, a dramatic reverse effect on oscillations was seen, the amplitude almost completely collapsing (Fig. S3).

**Fig. 5:**
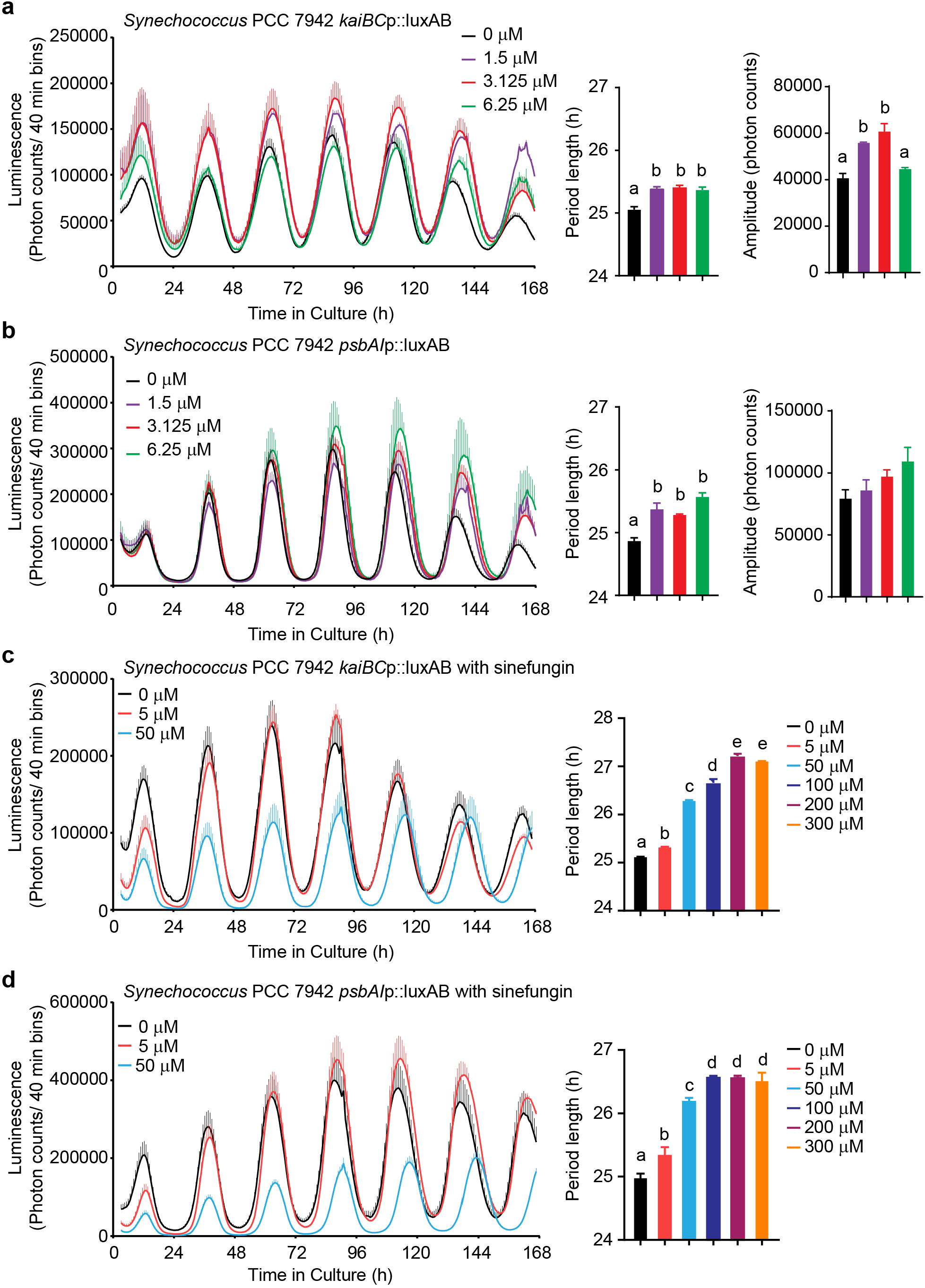
The link between the methyl cycle and the clock is conserved in cyanobacteria. **(a)** Left panel shows mean luminescence +/- SEM of *Synechococcus* PCC 7942 *kaiBC*p::luxAB knock-in strain treated with different concentrations of DZnep, n = 3, with only the upper section of the error bars shown for clarity. Middle panel shows mean period +/- SEM, n = 3. Right panel shows mean amplitude +/- SEM, n = 3. **(b)** Same as (a) but using *Synechococcus* PCC 7942 *psbAI*p::luxAB knock-in strain. Data for (a) and (b) were analyzed together with data from Fig. S3, showing additional concentrations of DZnep. **(c)** and **(d)** are the same as (a) and (b), respectively, but with different Sinefungin concentrations as indicated over the graphs. In addition, bar charts in (c) and (d) show data obtained with higher concentrations of Sinefungin presented in Fig. S3. See also Fig. S4 for a comparison with EGX^I^. All bar graphs analyzed by One-Way ANOVA followed by Bonferroni post-hoc test; all indicated comparisons at least *p* < 0.05.

Inhibition of methyltransferases by DZnep depends on the unique activity of AHCY to hydrolyze SAH. Some bacterial proteins, such as Mtn in *E.coli* or Pfu in *Streptococcus pyogenes*, however possess a SAH nucleosidase activity that cleaves SAH to adenine and ribosylhomocysteine (43). If this pathway for AHCY catabolism is active in *Synechococcus elongatus*, acting as a buffer against SAH accumulation, it would explain the blunted period response to DZnep compared to eukaryotes. Indeed, SAH nucleosidases have been identified in at least some *Synechococcus* strains, such as PCC7336 and MED-G69 (44). To circumvent the potential activity of a SAH nucleosidase in *Synechococcus elongatus*, we decided to use the global methylation inhibitor sinefungin, a natural analogue of SAM that directly binds to and inhibit methyltransferases (45, 46). A known antifungal and antibacterial agent whose activity as such has been shown to depend at least partially on the inhibition of mRNA 5’-cap methylation (47), it was previously used at 10 and 100 μM on the cyanobacteria *Anabaena* to elicit non-lethal methylation-dependent morphological changes (48). We thus selected Sinefungin to further probe the link between methyltransferases and the circadian period in *Synechococcus elongatus*. In both reporter strains, significant dose-dependent period lengthening was observed. To contrast with Sinefungin, we also tested a selective bacterial DNA methyltransferase inhibitor with a different molecular footprint from DZnep or Sinefungin, the cyclopentaquinoline carboxylic acid EGX^I^ (8-ethoxy-6-nitro-3a,4,5,9b-tetrahydro-3H- cyclopenta[c]quinoline-4-carboxylic acid) (49) (Fig. S4). Since its published IC_50_ for *E. coli* DNA methyltransferase is 9.7 μM (49), we used EGX^I^ at 5 and 50 μM but did not observe any consistent effects on the period, with only a mild period shortening in *kaiBC*p::luxAB cells at 50 μM (Fig S4). At 100 μM or higher, Sinefungin showed some growth inhibition during the experiment, but *Synechococcus* still showed healthy rhythms. In contrast, at 100 μM or higher EGX^I^ was very toxic to cyanobacteria, causing a dramatic collapse of luciferase intensity, with poor rhythms from which circadian parameters could not be reliably extracted.

Together these data show that the circadian clock in cyanobacteria —separated by 2 billion years of evolution from humans— is sensitive to methylation inhibition. This is especially meaningful considering that the cyanobacterial core clock is a phosphorylation-based biochemical oscillator. This also underlines the fundamental role that methyl metabolism kept in the control of physiology and behavior across evolution. Alternative metabolic pathways branching from the methyl cycle may prevent the accumulation of SAH and make prokaryotes more resistant to methylation inhibition by nutrient deprivation.

### The somite segmentation clock and methyl cycle are linked

Our results so far have shown that the link between the methyl cycle and the circadian TTFL is conserved. What about other biological timekeepers? While the circadian clock involves a TTFL that oscillates with a period near 24 hours, another TTFL, called the somite segmentation clock, cycles much faster and orchestrates the appearance of new somites from the paraxial mesoderm of the developing embryo in vertebrates (Fig. 6). In mouse, the underlying molecular oscillator is centered on the transcription factor *Hairy & Enhancer Of Split 7* (*Hes7*), whose expression oscillates with a period close to 2 hours by negative feedback that also induces oscillations of *Notch* and *Fgf* signaling (50, 51).

**Fig. 6:**
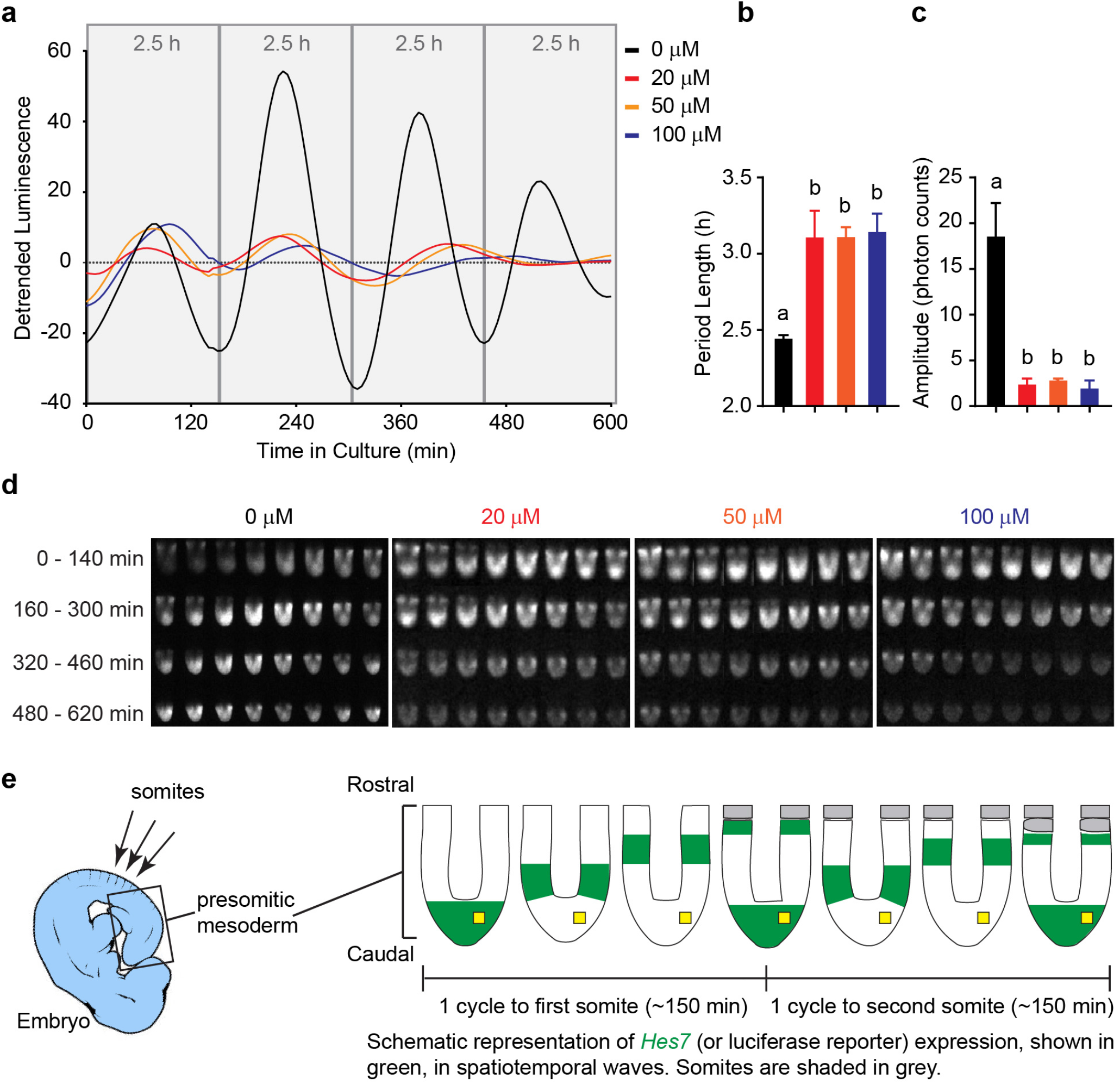
The somite segmentation clock is sensitive to methyl cycle inhibition. **(a)** Representative detrended luminescence measurements from one cultured embryo per treatment as indicated in the legend on the right. **(b)** Mean period +/- SEM of n = 3 embryos per treatment, with * = *p* < 0.05. **(c)** Mean amplitude +/- SEM of n = 3 embryos per treatment, with ** = *p* < 0.01. **(d)** Montage of luminescence time-lapse micrographs of one representative embryo for each treatment. One picture every 20 min is shown, starting from top left, each row corresponding to one *Hes7* expression cycle in the embryo treated with 0 μM DZnep, as shown in (a). See also Movies S2 and S3. **(e)** Anatomic localization of the presomitic mesoderm in embryo and schematic representation of the pictures from (d). The yellow square defined the area from which luminescence was measured in (a).

Like circadian clock genes, *Hes7* oscillatory expression can be monitored in real-time from transgenic embryo expressing highly destabilized luciferase under the control of the *Hes7* promoter. Testing increasing concentrations of DZnep on these transgenic embryos revealed uncanny similarities with results seen so far: an increase in period length accompanied by a decrease in amplitude of *Hes7* oscillations (Fig. 6). This once more illustrates the importance of the methyl cycle in the regulation of transcriptional programs orchestrating development, physiology and behaviour.

Here we provide evidence for the existence of an ancient link between the methyl cycle and biological timekeepers. In the course of more than 2 billion years of cellular evolution, during which the biological clock and the methyl cycle have adapted to a multicellular eukaryotic existence, the link between these two processes has remained active. Further investigations should reveal how the link itself has evolved, and which methylated substrates and methyltransferases are critical for its function.

## DISCUSSION

Solar ultraviolet radiation played a critical role in prebiotic chemistry and has been involved in the origin of chirality (*52*), the synthesis of alcohols, aldehydes (notably formaldehyde), and organic acids (*53*), as well as that of amino acids (*54*) and ribonucleotides (*55*). The presence of methylation transfer in the prebiotic world probably caused the incorporation of methyl chemistry in early life-forms. The energy of UV photons however can also degrade biologically important molecules, preventing abiogenesis. Once autocatalytic pseudo-life forms evolved with more complex chemistry, avoidance mechanisms may have provided increased fitness in such an environment. The origin of the circadian clock on the early Earth is likely to have been a simple sensing of molecules present in the milieu, increasing or decreasing under UV exposure and triggering an appropriate response. Formaldehyde, increasing during the day under the action of UV on carbon monoxide and water vapor (*53*), could have been such a chemical marker of daylight, affecting the chemistry of early life-forms by methylation. A conserved link between methyl transfer and the circadian clock may have arisen from such a scenario.

The most surprising data obtained here is the almost identical response of unicellular algae and vertebrates to DZnep despite their evolutionary divergence more than 1 billion years ago, the comparable effect of methylation inhibitors on cyanobacterial circadian clock (2 billion years of evolutionary divergence), and the conservation of this biochemical link in the control of the somite segmentation clock. So far, only a small number of examples have been reported that identified aspects of circadian timekeeping shared among rhythmic life, such as oxidation cycles of peroxiredoxin (34, 56, 57) and regulation by oscillating intracellular magnesium concentrations (58).

In mammalian cells we previously reported that methyl cycle inhibition decreased the methylation of internal adenosines in mRNA (m6A), as well as that of histones (*8*). While specific m6A inhibition was sufficient to elicit period elongation, the contribution of histone methylation to the period elongation obtained by DZnep was likely significant, as well as that of other methylation sites in mRNA, rRNA and tRNA. Due to the considerable heterogeneity in mechanisms regulating gene expression and function in organisms tested here, identifying a single mechanism explaining period elongation would be a difficult undertaking and is beyond the scope of the present work. We mentioned in the introduction that the oldest MTases are RNA methyltransferases (*5*), it is therefore tempting to propose that inhibition of RNA methylation may at least partially contribute to the period lengthening. That sinefungin, a confirmed mRNA cap-methyltransferase inhibitor (47), was able to lengthen the cyanobacterial clock period while EGX^I^, a DNA methylation inhibitor, was not support this hypothesis. It is also possible that the effects of methyl cycle inhibition on period and amplitude in eukaryotes arise from different mechanisms, e.g. on the inhibition of mRNA and histone methylation, respectively. More experiments should clarify these points.

The presence of a SAH nucleosidase in *S. elongatus* may have blunted the effect of DZnep, but even with 100 μM sinefungin the period lengthened only about 2 hours. It should be mentioned that our luciferase reporter system is a reporter for the driven TTFL, not for the KaiABC oscillator *per se*. It is therefore possible that methylation inhibition only affected the coupling between the nanocomplex and the TTFL, which may explain why the increase in period was less pronounced compared to most eukaryotes tested here. In eukaryotic cells, luciferase reporter systems are a direct read-out of the core oscillator that has evolved more dependent on the TTFL, and as a result may have become more sensitive to perturbations affecting gene expression such as inhibition of RNA and histone methylation.

The somite segmentation clock was strongly affected by inhibition of AHCY, which is in line with the importance of 1 carbon metabolism for embryonic development. As can be seen in the Movie S3, showing similar results to Movie S2 but merged with brightfield images and displaying luminescence as a pseudo-color green, period lengthening of the oscillatory expression of *Hes7-luciferase* occurred together with a pronounced delay in the growth of the caudal tip of the presomitic mesoderm as well as in the appearance of new somites, demonstrating that the molecular clock as well as its output, i.e. the somitogenesis, were affected by DZnep. The mechanisms underlying this period lengthening may be distinct from those involved in the lengthening of the circadian period. Considering the short period of *Hes7* oscillations — 2 h, and the importance of 3’-UTR-dependent regulation of *Hes7* mRNA turnover for its cyclic expression (62), the inhibition of mRNA methylation may at least in part contribute to the results observed.

Despite being one of the most potent AHCY inhibitors (11), DZnep is sometimes erroneously sold and used as a “specific” histone methyltransferase EZH2 inhibitor because of the misinterpretation of a report showing that it inhibits —without any data on specificity— histone methylation by EZH2 in cancer cells (59). More recent reports showed that, in line with its inhibitory effect on AHCY (60, 61), DZnep globally inhibits histone methylation.

Since we sought to inhibit the methyl cycle in organisms from bacteria to humans, these investigations were by necessity limited to the pharmacological inhibition of AHCY, because irreversible genetic disruption of AHCY in all organisms would have been much less feasible, notably due to likely embryonic lethality or developmental arrest in metazoa, cell cycle/growth disruption, and the existence of multiple uncharacterized homologues in many organisms tested here. This work puts the spotlight on the methyl cycle and calls for more investigations into how it regulates physiology and behavior.

## AKNOWLEDGEMENTS

This work was supported in part by the Ministry of Education, Culture, Sports, Science and Technology of Japan: Grant-in-aid for Scientific Research on Innovative Areas (26116713, J.-M. F.), for Young Scientists (26870283, J.-M. F.), for Scientific Research A (15H01843, H.O.), a grant for Core Research for Evolutional Science and Technology, Japan Science and Technology Agency (CREST/JPMJCR14W3, H.O.). J.-M.F. was also supported by grants from the Kato Memorial Bioscience Foundation, the Senri Life Science Foundation (S-26003), the Mochida Memorial Foundation for Medical and Pharmaceutical Research, and the Kyoto University internal grant ISHIZUE, and H.O. was also supported by the Kobayashi International Scholarship Foundation. C.H.J. is supported by grants from the USA NIH/NIGMS: GM067152 and GM107434. G.vO. is supported by a Royal Society University Research Fellowship (UF160685) and research grant (RGF\EA\180192). D.A.G. is supported by grants from the National Research Agency (ANPCyT, PICT-2015-0572) and the National University of Quilmes. We thank Adrienne K. Mehalow, Jay C. Dunlap, John O’Neill and Tokitaka Oyama for useful discussion and for kindly accepting to perform experiments. The authors thank G. Wolber, Freie Universität Berlin, Germany, for providing a LigandScout 4.2 license.

## Author Contributions

Conceptualization, J.-M.F. and H.O.; Methodology, J.-M.F., C.R., M.V., S.J.C., T.K.T., Y.X., M.L.J., M.L.L., K.Y.-K., D.W., R.Ka., T.M., R.S., D.A.G., C.H.J., G.v.O.; investigation, J.-M.F., S.Y., C.R., M.V., S.J.C., T.K.T., Y.X., M.L.J., R.Ko., M.L.L., K.Y.-K., T.M., D.A.G., G.v.O.; homology modelling, C.R.; original draft, J.-M.F.; reviewing & editing, J.-M.F., C.R., T.K.T, Y.X., K.Y.-K., D.W., R.Ka., T.M., R.S., D.A.G., C.H.J., G.v.O.; Funding Acquisition, J.-M.F., D.A.G., C.H.J., G.v.O and H.O.; Supervision, J.-M.F.

## METHODS

### Molecular modelling and sequence analyses

Homology models were obtained as described in the SI Appendix.

### Assay for effect of the inhibition of the methyl cycle on vertebrate circadian clock

Human U-2 OS cells stably transfected with a *Bmal1*-luciferase reporter vector (22) and mouse PER2::LUC MEFs (23) cell lines were cultivated as previously described (8). Briefly, cells were seeded into 35 mm dishes (Corning) and allow to grow to confluence in DMEM/F12 medium (Nacalai) containing penicillin/streptomycine/amphotericin (Nacalai). Cells were then shocked by dexamethasone (Sigma-Aldrich) 200 nM for 2 hours, followed by a medium changed including 1 mM luciferine (Nacalai). 35 mm dishes were then transferred to an 8-dishes luminometer-incubator (Kronos Dio, Atto). Photons were counted in bins of 2 min at a frequency of 10 min. DZnep was purchased from Sigma-Aldrich.

Zebrafish cell line and bioluminescence assays. The generation of a per1b-luciferase cell line from zebrafish PAC2 cells has been previously described (*64*). Cells were cultured in Leibovitz’s L-15 medium (Gibco) containing 15% fetal bovine serum (Biochrom AG), 50 U/mL penicillin/streptomycin (Gibco), and 50 μg/mL gentamicin (Gibco). Cells were seeded at a density of 50,000-100,000 cells per well in quadruplicate wells of a 96-well plate in medium supplemented with 0.5 mM beetle luciferin (Promega), and drugs were prepared in water and added at the concentrations indicated in the figures and legends. Plates were sealed with clear adhesive TopSeal (Perkin Elmer, Waltham, MA, USA). Cells were exposed to a 12:12 LD cycle for 7 days and then transferred into DD for at least 3 days. Bioluminescence was monitored on a Packard TopCount NXT scintillation counter (28°C). DZnep was purchased from Sigma-Aldrich.

Period and amplitude were estimated by BioDare2 (63).

### Assay for effect of the inhibition of the methyl cycle on invertebrate circadian clock

Halteres of two-to seven-day old transgenic *ptim*-TIM-LUC males (65) kept under 12 h :12 h light:dark cycles (LD) at 25°C were bilaterally dry dissected. Each pair was transferred into one well of a 96 well plate (Topcount, Perkin Elmer) filled with medium containing 80% Schneider’s medium (Sigma), 20% inactivated Fetal Bovine Serum (Capricorn) and 1% PenStrep (Sigma). Medium was fortified with 226 µM Luciferin (Biosynth) and supplemented with 10, 100, or 250 µM Dznep A diluted in PBS. Plates were sealed with clear adhesive covers and transferred to a TopCount plate reader (PerkinElmer). Bioluminescence emanating from each well was measured hourly in LD for two days, followed by 5 days of constant darkness (DD) at 25°C as previously described (66). The *ptim*-TIM-LUC reporter contains the timeless (*tim*) promoter sequences driving rhythmic expression of the *tim* cDNA, which is fused to the firefly luciferase cDNA.

*C*. *elegans* strain N2 (Bristol strain, wild-type was provided by the Caenorhabditis Genetics Center, University of Minnesota (cbs.umn.edu/cgc/home). Stocks were maintained on plates with nematode growth medium (NGM) seeded with HB101 *Escherichia coli* strain, under 12-h/12-h LD/CW cycle (400/0 lx and CW (18.5/20 °C, Δ = 1.5 ± 0.125 °C) environmental cycles. Transgenic animals were generated by microinjection of a P*sur-5::luc::gfp* construct at 50 or 100 ng/μL with the pRF4 marker (100 ng/μL) (*29, 67*). Bioluminescence recordings with nematodes are described in the SI Appendix.

Period and amplitude were estimated by BioDare2 (63).

### Assay for effect of the inhibition of the methyl cycle on plant and algae circadian clock

*Ostreococcus tauri* cells transgenically expressing a translational fusion of CCA1 to luciferase from the CCA1 promoter (CCA1-LUC) (*36*) were grown, imaged, and analysed as described previously (*58*).

For protoplast isolation and luminescent imaging, plants expressing luciferase from the CCA1 promoter in the Col-0 background (kindly provided by Karen Halliday, University of Edinburgh) were grown in sterile soil under long-day conditions (16 h light/8 h dark) at 22°C under 60-70 µmol m^−2^s^−1^ white LED tube lights (Impact T8). Protoplasts were isolated from 3-week old leaves as previously described (68). Protoplasts in a solution of 1 mM D-luciferin (Biosynth AG), 5 % fetal bovine serum (Sigma), 50µg/ml ampicillin, 140 mM NaCl, 115 mM CaCl2, 4.6 mM KCl, 1.86 mM MES pH 5.7 and 4.6 mM glucose were added to white, flat-bottomed 96-well plates (Lumitrac, Greiner Bio-one) at a concentration of 2×10^5^ cells/ml. DZnep (S7120 Selleckchem) was added to the protoplasts to achieve concentrations of 0, 0.125, 0.25, 0.5 or 1 µM from a 1 mM stock solution (prepared in distilled H_2_O) to a total volume of 200 µL per well. Plates were sealed with a clear adhesive lid (TopSeal-A, Perkin Elmer). Protoplast luminescence was read by a LB942 Tristar^2^ plate reader (Berthold Technologies Ltd) every 50 minutes for 3 seconds per well, and kept under continuous red (630 nm) and blue (470 nm) LED light (5 µmol m^−2^s^−1^ each) at 19°C.

*Chlamydomonas reinhardtii* strain CBR carrying a codon-adapted luciferase reporter driven by the *tufA* promoter in the chloroplast genome (69, 70) was used. Culture preparation, bioluminescence monitoring, and data analysis were carried out as described previously (70). Briefly, 5-day-old algal cultures on HS agar medium were cut out along with the agar by using a glass tube, and transferred to separate wells of a 96-well microtiter plates. Luciferin (final conc. 200 μM) and various concentrations of DZnep (Sigma) were added to the wells. Algae were synchronized by a single cycle of 12-h darkness/12-h light (30 μmol m^−2^ s^−1^) at 17°C before bioluminescence monitoring using a custom-made luminometer-incubator (71) in DD at 17°C.

Period and amplitude were estimated by BioDare2 (63).

### Assay for effect of the inhibition of the methyl cycle on prokaryotic circadian clock

*Vibrio harveyi* luciferase encoded by *luxA* and *luxB* (*luxAB*) genes was used as a luminescence reporter in cyanobacterium *Synechococcus elongatus* PCC 7942. Two cyanobacterial clock-regulated reporter strains, *kaiBC*p::*luxAB* (*72*) and *psbAI*p::*luxAB* (*73*) were selected for the test, *i.e.* expression of the *luxAB* was under control of the promoters of the central clock genes *kaiBC* and the Class I photosynthetic gene *psbAI*, respectively. *Synechococcus* strains were grown in modified BG11 media (*74*) supplemented with 40 µg/ml of spectinomycin. The cultures grown on BG11 agar plates for 3 days at 30°C under continuous cool-white illumination (LL) (50 µE/m^2^ s) were toothpicked onto fresh BG11 agar plates containing 0.015 g/L of L-methionine and different concentrations of 3-deazaneplanocin A hydrochloride (Sigma), sinefungin (Abcam) or EGX^I^ (ChemBridge 5790780). After a single 12h dark pulse for synchronization, assay of the *in vivo* luminescence rhythms was performed as described previously (*72*).

Period and amplitude were estimated by BioDare2 (63).

### Measurement of *Hes7* oscillations in the mouse presomitic mesoderm

Presomitic mesoderm (PSM) tissues from three littermate p*Hes7*-UbLuc Tg embryos were embedded in 0.35% LMP-agarose/culture medium (10% FBS-DMEM/F12), in a silicon mold mounted onto a φ35mm-glass bottom dish, and luciferin-containing medium with DZnep or vehicle (MilliQ water) were added.

Time-lapse imaging of PSM was performed with an inverted microscope (Olympus IX81) equipped with an Olympus x10 UPlanApo objective (N.A.: 0.8) and a VersArray cooled-CCD camera. 16-bit images were acquired every 5min with Image-Pro Plus (Media Cybernetics), with 2×2 and 4×4 binning and exposures of 100 ms and 4 m 25 s for DIC and chemiluminescent images, respectively. Raw imaging data were processed and quantified by ImageJ (75).

Period and amplitude were estimated by BioDare2 (63).

## References

1. Lippi G & Plebani M (2012) Hyperhomocysteinemia in health and disease: Where we are now, and where do we go from here ?. Clin Chem Lab Med 50(12): 2075–2080.

2. Cantoni GL (1985) The role of S-adenosylhomocysteine in the biological utilization of S-adenosylmethionine. Prog Clin Biol Res 198: 47–65.

3. Chiang PK & Cantoni GL (1979) Perturbation of biochemical transmethylations by 3-deazaadenosine in vivo. Biochem Pharmacol 28(12): 1897–1902.

4. Hoffman DR, Marion DW, Cornatzer WE & Duerre JA (1980) S-adenosylmethionine and S-adenosylhomocystein metabolism in isolated rat liver. effects of L-methionine, L-homocystein, and adenosine. J Biol Chem 255(22): 10822–10827.

5. Rana AK & Ankri S (2016) Reviving the RNA world: An insight into the appearance of RNA methyltransferases. Front Genet 7: 99.

6. Van Trump JE & Miller SL (1972) Prebiotic synthesis of methionine. Science 178(4063): 859–860.

7. Waddell TG, Eilders LL, Patel BP & Sims M (2000) Prebiotic methylation and the evolution of methyl transfer reactions in living cells. Orig Life Evol Biosph 30(6): 539–548.

8. Fustin JM, et al (2013) RNA-methylation-dependent RNA processing controls the speed of the circadian clock. Cell 155(4): 793–806.

9. Guranowski A, Montgomery JA, Cantoni GL & Chiang PK (1981) Adenosine analogues as substrates and inhibitors of S-adenosylhomocysteine hydrolase. Biochemistry 20(1): 110–115.

10. Chiang PK, Richards HH & Cantoni GL (1977) S-adenosyl-L-homocysteine hydrolase: Analogues of S-adenosyl-L-homocysteine as potential inhibitors. Mol Pharmacol 13(5): 939–947.

11. Tseng CK, et al (1989) Synthesis of 3-deazaneplanocin A, a powerful inhibitor of S-adenosylhomocysteine hydrolase with potent and selective in vitro and in vivo antiviral activities. J Med Chem 32(7): 1442–1446.

12. Keiser MJ, et al (2007) Relating protein pharmacology by ligand chemistry. Nat Biotechnol 25(2): 197–206.

13. Gfeller D, Michielin O & Zoete V (2013) Shaping the interaction landscape of bioactive molecules. Bioinformatics 29(23): 3073–3079.

14. Yang X, et al (2003) Catalytic strategy of S-adenosyl-L-homocysteine hydrolase: Transition-state stabilization and the avoidance of abortive reactions. Biochemistry 42(7): 1900–1909.

15. Kusakabe Y, et al (2015) Structural insights into the reaction mechanism of S-adenosyl-L-homocysteine hydrolase. Sci Rep 5: 16641.

16. Brzezinski K, Dauter Z & Jaskolski M (2012) High-resolution structures of complexes of plant S-adenosyl-L-homocysteine hydrolase (lupinus luteus). Acta Crystallogr D Biol Crystallogr 68(Pt 3): 218–231.

17. Gomi T, et al (1990) Site-directed mutagenesis of rat liver S-adenosylhomocysteinase. effect of conversion of aspartic acid 244 to glutamic acid on coenzyme binding. J Biol Chem 265(27): 16102–16107.

18. Ault-Riche DB, Yuan CS & Borchardt RT (1994) A single mutation at lysine 426 of human placental S-adenosylhomocysteine hydrolase inactivates the enzyme. J Biol Chem 269(50): 31472–31478.

19. Takata Y, et al (2002) Catalytic mechanism of S-adenosylhomocysteine hydrolase. site-directed mutagenesis of asp-130, lys-185, asp-189, and asn-190. J Biol Chem 277(25): 22670–22676.

20. Yamada T, et al (2005) Catalytic mechanism of S-adenosylhomocysteine hydrolase: Roles of his 54, Asp130, Glu155, Lys185, and Aspl89. Int J Biochem Cell Biol 37(11): 2417–2435.

21. Sganga MW, Aksamit RR, Cantoni GL & Bauer CE (1992) Mutational and nucleotide sequence analysis of S-adenosyl-L-homocysteine hydrolase from rhodobacter capsulatus. Proc Natl Acad Sci U S A 89(14): 6328–6332.

22. Baggs JE, et al (2009) Network features of the mammalian circadian clock. PLoS Biol 7(3): e52.

23. Yoo SH, et al (2004) PERIOD2::LUCIFERASE real-time reporting of circadian dynamics reveals persistent circadian oscillations in mouse peripheral tissues. Proc Natl Acad Sci U S A 101(15): 5339–5346.

24. Bargiello TA, Jackson FR & Young MW (1984) Restoration of circadian behavioural rhythms by gene transfer in drosophila. Nature 312(5996): 752–754.

25. Young MW, Jackson FR, Shin HS & Bargiello TA (1985) A biological clock in drosophila. Cold Spring Harb Symp Quant Biol 50: 865–875.

26. Hardin PE, Hall JC & Rosbash M (1990) Feedback of the drosophila period gene product on circadian cycling of its messenger RNA levels. Nature 343(6258): 536–540.

27. Kippert F, Saunders DS & Blaxter ML (2002) Caenorhabditis elegans has a circadian clock. Curr Biol 12(2): R47–9.

28. Saigusa T, et al (2002) Circadian behavioural rhythm in caenorhabditis elegans. Curr Biol 12(2): R46–7.

29. Goya ME, Romanowski A, Caldart CS, Benard CY & Golombek DA (2016) Circadian rhythms identified in caenorhabditis elegans by in vivo long-term monitoring of a bioluminescent reporter. Proc Natl Acad Sci U S A 113(48): E7837–E7845.

30. Lee S, Doxey AC, McConkey BJ & Moffatt BA (2012) Nuclear targeting of methyl-recycling enzymes in arabidopsis thaliana is mediated by specific protein interactions. Mol Plant 5(1): 231–248.

31. Grbesa I, et al (2017) Mutations in S-adenosylhomocysteine hydrolase (AHCY) affect its nucleocytoplasmic distribution and capability to interact with S-adenosylhomocysteine hydrolase-like 1 protein. Eur J Cell Biol 96(6): 579–590.

32. Matsuo T & Ishiura M (2011) Chlamydomonas reinhardtii as a new model system for studying the molecular basis of the circadian clock. FEBS Lett 585(10): 1495–1502.

33. van Ooijen G, et al (2013) Functional analysis of casein kinase 1 in a minimal circadian system. PLoS One 8(7): e70021.

34. O’Neill JS, et al (2011) Circadian rhythms persist without transcription in a eukaryote. Nature 469(7331): 554–558.

35. van Ooijen G, Dixon LE, Troein C & Millar AJ (2011) Proteasome function is required for biological timing throughout the twenty-four hour cycle. Curr Biol 21(10): 869–875.

36. Corellou F, et al (2009) Clocks in the green lineage: Comparative functional analysis of the circadian architecture of the picoeukaryote ostreococcus. Plant Cell 21(11): 3436–3449.

37. Sassone-Corsi P & Christen Y (2016) A Time for Metabolism and Hormones, (Springer International Publishing : Imprint: Springer, Cham), pp 132.

38. Nakajima M, et al (2005) Reconstitution of circadian oscillation of cyanobacterial KaiC phosphorylation in vitro. Science 308(5720): 414–415.

39. Johnson CH, Zhao C, Xu Y & Mori T (2017) Timing the day: What makes bacterial clocks tick?. Nat Rev Microbiol 15(4): 232–242.

40. Ishiura M, et al (1998) Expression of a gene cluster kaiABC as a circadian feedback process in cyanobacteria. Science 281(5382): 1519–1523.

41. Liu Y, et al (1995) Circadian orchestration of gene expression in cyanobacteria. Genes Dev 9(12): 1469–1478.

42. Qin X, Byrne M, Xu Y, Mori T & Johnson CH (2010) Coupling of a core post-translational pacemaker to a slave transcription/translation feedback loop in a circadian system. PLoS Biol 8(6): e1000394.

43. Walker RD & Duerre JA (1975) S-adenosylhomocysteine metabolism in various species. Can J Biochem 53(3): 312–319.

44. Shih PM, et al (2013) Improving the coverage of the cyanobacterial phylum using diversity-driven genome sequencing. Proc Natl Acad Sci U S A 110(3): 1053–1058.

45. Hamill RL & Hoehn MM (1973) A9145, a new adenine-containing antifungal antibiotic. J Antibiot (Tokyo) 26: 463.

46. Vedel M, Lawrence F, Robert-Gero M & Lederer E (1978) The antifungal antibiotic sinefungin as a very active inhibitor of methyltransferases and of the transformation of chick embryo fibroblasts by rous sarcoma virus. Biochem Biophys Res Commun 85(1): 371–376.

47. Zheng S, et al (2006) Mutational analysis of encephalitozoon cuniculi mRNA cap (guanine-N7) methyltransferase, structure of the enzyme bound to sinefungin, and evidence that cap methyltransferase is the target of sinefungin’s antifungal activity. J Biol Chem 281(47): 35904–35913.

48. Fujita H, Syono K, Machida Y & Kawaguchi M (2008) Morphological effects of sinefungin, an inhibitor of S-adenosylmethionine-dependent methyltransferases, on anabaena sp. PCC 7120. Microbes Environ 23(4): 346–349.

49. Mashhoon N, Pruss C, Carroll M, Johnson PH & Reich NO (2006) Selective inhibitors of bacterial DNA adenine methyltransferases. J Biomol Screen 11(5): 497–510.

50. Kageyama R, Niwa Y, Isomura A, Gonzalez A & Harima Y (2012) Oscillatory gene expression and somitogenesis. Wiley Interdiscip Rev Dev Biol 1(5): 629–641.

51. Kobayashi T & Kageyama R (2014) Expression dynamics and functions of hes factors in development and diseases. Curr Top Dev Biol 110: 263–283.

52. Rosenberg RA, Abu Haija M & Ryan PJ (2008) Chiral-selective chemistry induced by spin-polarized secondary electrons from a magnetic substrate. Phys Rev Lett 101(17): 178301.

53. Bar-Nun A & Hartman H (1978) Synthesis of organic compounds from carbon monoxide and water by UV photolysis. Orig Life 9(2): 93–101.

54. Sarker PK, Takahashi J, Obayashi Y, Kaneko T & Kobayashi K (2013) Photo-alteration of hydantoins against UV light and its relevance to prebiotic chemistry. Advances in Space Research 51(12): 2235–2240.

55. Powner MW, Gerland B & Sutherland JD (2009) Synthesis of activated pyrimidine ribonucleotides in prebiotically plausible conditions. Nature 459(7244): 239–242.

56. O’Neill JS & Reddy AB (2011) Circadian clocks in human red blood cells. Nature 469(7331): 498–503.

57. Edgar RS, et al (2012) Peroxiredoxins are conserved markers of circadian rhythms. Nature 485(7399): 459–464.

58. Feeney KA, et al (2016) Daily magnesium fluxes regulate cellular timekeeping and energy balance. Nature 532(7599): 375–379.

59. Tan J, et al (2007) Pharmacologic disruption of polycomb-repressive complex 2-mediated gene repression selectively induces apoptosis in cancer cells. Genes Dev 21(9): 1050–1063.

60. Miranda TB, et al (2009) DZNep is a global histone methylation inhibitor that reactivates developmental genes not silenced by DNA methylation. Mol Cancer Ther 8(6): 1579–1588.

61. Zeybel M, et al (2017) A proof-of-concept for epigenetic therapy of tissue fibrosis: Inhibition of liver fibrosis progression by 3-deazaneplanocin A. Mol Ther 25(1): 218–231.

62. Nitanda Y, et al (2014) 3’-UTR-dependent regulation of mRNA turnover is critical for differential distribution patterns of cyclic gene mRNAs. Febs J 281(1): 146–156.

63. Zielinski T, Moore AM, Troup E, Halliday KJ & Millar AJ (2014) Strengths and limitations of period estimation methods for circadian data. PLoS One 9(5): e96462.

64. Vallone D, Gondi SB, Whitmore D & Foulkes NS (2004) E-box function in a period gene repressed by light. Proc Natl Acad Sci U S A 101(12): 4106–4111.

65. Lamba P, Foley LE & Emery P (2018) Neural network interactions modulate CRY-dependent photoresponses in drosophila. J Neurosci 38(27): 6161–6171.

66. Chen C, Xu M, Anantaprakorn Y, Rosing M & Stanewsky R (2018) Nocte is required for integrating light and temperature inputs in circadian clock neurons of drosophila. Curr Biol 28(10): 1595–1605.e3.

67. Mello CC, Kramer JM, Stinchcomb D & Ambros V (1991) Efficient gene transfer in C.elegans: Extrachromosomal maintenance and integration of transforming sequences. Embo J 10(12): 3959–3970.

68. Hansen LL & van Ooijen G (2016) Rapid analysis of circadian phenotypes in arabidopsis protoplasts transfected with a luminescent clock reporter. J Vis Exp (115). doi(115): 10.3791/54586.

69. Hwang S, Kawazoe R & Herrin DL (1996) Transcription of tufA and other chloroplast-encoded genes is controlled by a circadian clock in chlamydomonas. Proc Natl Acad Sci U S A 93(3): 996–1000.

70. Matsuo T, et al (2008) A systematic forward genetic analysis identified components of the chlamydomonas circadian system. Genes Dev 22(7): 918–930.

71. Okamoto K, Onai K, Ezaki N, Ofuchi T & Ishiura M (2005) An automated apparatus for the real-time monitoring of bioluminescence in plants. Anal Biochem 340(2): 187–192.

72. Xu Y, Mori T & Johnson CH (2003) Cyanobacterial circadian clockwork: Roles of KaiA, KaiB and the kaiBC promoter in regulating KaiC. Embo J 22(9): 2117–2126.

73. Kondo T, et al (1993) Circadian rhythms in prokaryotes: Luciferase as a reporter of circadian gene expression in cyanobacteria. Proc Natl Acad Sci U S A 90(12): 5672–5676.

74. Bustos SA & Golden SS (1991) Expression of the psbDII gene in synechococcus sp. strain PCC 7942 requires sequences downstream of the transcription start site. J Bacteriol 173(23): 7525–7533.

75. Shimojo H, Harima Y & Kageyama R (2014) Visualization of notch signaling oscillation in cells and tissues. Methods Mol Biol 1187: 169–179.

