## Supplemental methods and data for "The methyl cycle is a conserved regulator of biological clocks"

**This PDF file includes:**

Supplementary text
Figs. S1 to S4
Captions for movies S1 to S3

**Other supplementary materials for this manuscript include the following:**

Movies S1 to S3

### Supplementary Methods

#### Molecular modelling and sequence analyses

3D protein structures were obtained from the Protein Data Bank (PDB) for *Homo sapiens* (PDB 1LI4 (1) , S-adenosylhomocysteine hydrolase complexed with neplanocin, resolution 2.01 Å) and *Mus musculus* (PDB 5AXA (2), S-adenosylhomocysteine hydrolase with adenosine, chain A, resolution 1.55 Å). All PDB structures underwent preprocessing including the addition of missing amino acid atoms and hydrogens, removal of solvent atoms, and hydrogen-bond network optimization and residue protonation based on predicted pKa values using the Protein Preparation Wizard (3) of Maestro Schrödinger Release 2018-1 [Maestro, Schrödinger, LLC, New York, NY, 2018]. Homology models of *Danio rerio* (template 1LI4, 100 % coverage, 86 % sequence identity), *Drosophila melanogaster* (template 1LI4, 100 % coverage, 81 % sequence identity), *Caenorhabditis elegans* (template 1LI4, 98 % coverage, 77 % sequence identity), *Arabidopsis thaliana* (template 3OND (4) chain A, resolution 1.17 Å, 100 % coverage, sequence 92 % identity), *Chlamydomonas reinhardtii* (template 3OND chain A, 100 % coverage, sequence 80 % identity), *Ostreococcus tauri* (template 3OND chain A, 99 % coverage, 79 % sequence identity), and *Synechococcus elongatus* (template 1LI4, 98 % coverage, 41 % sequence identity) were constructed via the SWISS-MODEL server (5). Model quality was evaluated based on GMQE (global model quality estimation) ranging from zero to one with high numbers indicating high model reliability, and QMEAN (6), a composite and absolute measure for the quality of protein models, indicating good or poor agreement between model and template with values of around 0 or below -4, respectively. GMQE and QMEAN values for the seven homology models were determined to be 0.98 and 0.85 for *D. rerio*, 0.93 and 0.82 for *D. melanogaster*, 0.87 and 0.35 for *C. elegans*, 0.99 and 0.31 for *A. thaliana*, 0.95 and 0.08 for *C. reinhardtii*, 0.91 and -0.54 for *O. tauri*, and 0.75 and -0.59 for *S. elongatus*, respectively. GROMACS version 2018 (7) was utilised for energy minimization of all nine protein structures including oxidised cofactor $NAD^+$ using force field AMBERff99SB-ILDN (8). Cofactor parameters and topologies were obtained through ACPYPE (9) using the AM1-BCC method with net charge $q = -1$. The systems including TIP3P water were neutralized with counter ions ($Cl^-$ and $Na^+$) to 0.1 M and minimized via steepest descent with a maximum force limit of 100 kJ $mol^{-1}$ $nm^{-1}$. Molecular docking simulations were performed using the energy-minimized protein-$NAD^+$ complexes as well as co-crystallised adenosine (ADN), neplanocin (NOC), and 3-deazaneplanocin A (DZnep, constructed from co-crystal NOC in 1LI4 (1)) using the Lamarckian Genetic Algorithm provided by the AutoDock4.2 suite (10). Protein-ligand interactions were analysed in LigandScout version 4.2 (11, 12).

Multiple sequence alignment of AHCY was performed by CLUSTALW (13) with the multiple sequence viewer from the Maestro module using the following reference sequences: for human, Genbank: NP_000678.1; mouse, Genbank: NP_057870.3; zebrafish, Genbank: NP_954688.1; fruit fly, Genbank: NP_511164.2; *C. elegans*, Genbank: NP_491955.1; *Arabidopsis*: NP_193130.1; *Chlamydomonas*, Genbank: XP_001693339.1; *Ostreococcus*, Genbank: XP_022839640.1; cyanobacteria, Genbank: WP_011243218.1.

For the molecular structure of drugs shown in Fig. S4, the Open Source web application MolView was used.

**Bioluminescence recordings with nematodes.**

For bioluminescence recordings, transgenic P*sur-5::luc::gfp* nematode populations were synchronized to the same developmental stage by the chlorine method *(14)*. The harvested eggs were cultured overnight in a 50-mL Erlenmeyer flask with 3.5 mL of M9 buffer, 1× antibiotic-antimycotic (Thermo Fisher Scientific), and 10 µg/mL of tobramycin (Tobrabiotic; Denver Farma) at 110 rpm with a Vicking M23 shaker, in LD/CW (400/0 lx; 18.5/20 °C, Δ = 1.5 °C ± 0.125 °C) conditions. The following day, L1 larvae were transferred to NGM plates at ZT1 and were grown for 48 h to the L4 stage under the same LD/CW cycle. Starting at ZT1 (1 h post lights on), the most fluorescent nematodes were selected manually under a SMZ100 stereomicroscope equipped with an epi-fluorescence attachment (Nikon) with a cool Multi-TK-LED light source (Tolket) to avoid warming the plate. Picking was performed in a room kept at a constant temperature (18 °C). L4 larvae were selected after 48 h of light-dark/temperature entrainment and placed in a luminescence medium containing Leibovitz's L-15 medium without phenol red (Thermo Fisher Scientific) supplemented with 1× antibiotic–antimycotic (Thermo

Fisher), 40 μM of 5-fluoro-2′-deoxyuridine (FUdR) to avoid new eclosions, 5 mg/mL cholesterol, 10 μg/mL tobramycin (Tobrabiotic; Denver Farma), 1 mM D-luciferin (Gold Biotechnology), and 0.05% Triton X-100 to increase cuticle permeabilization. All chemical compounds were purchased from Sigma-Aldrich unless otherwise specified. Population nematode bioluminescence was recorded as a measure of circadian locomotor activity in 35-mm plate dishes (Greiner CELLSTAR) with 1 mL of the luminescence medium, by means of an AB-2550 Kronos Dio luminometer (ATTO). The signal was integrated for 1 min, and readings were taken every 10 min.

For single-nematode measurements, P*sur-5::luc::gfp* nematodes at the L4 stage were selected as before, washed, and transferred directly to the liquid luminescence medium. A small, square piece of a transparent 96-well plate containing three × three wells was inserted inside a 35-mm dish plate to reduce the total volume and limit the nematode movement to the center of the plate. One nematode was placed in the center well with 200 μL of the luminescence medium. Plates were then transferred to the AB-2550 Kronos Dio luminometer. Single-nematode recordings were taken every 37 min with an integration time of 4 min.

In both cases, 2 μl of 10 mM DZNep (in PBS 1x, pH 7.2) were added to the luminescence medium (200 μl/well), with a final concentration of 100 μM. Controls were treated with vehicle (PBS).

As previously described, custom MATLAB scripts (provided in SI) were used to analyse the raw luminescence data (15). This is necessary because of the high signal/noise ratio, especially for single worms. Briefly, data were trend-corrected by dividing by a fixed moving-average window of 24h, then smoothed for 12h. Data were further normalized to the maximum level of luminescence of all biological replicates along the entire time series on each experiment. After raw data clean-up by MATLAB, period and amplitude were analysed by BioDare2 (16).

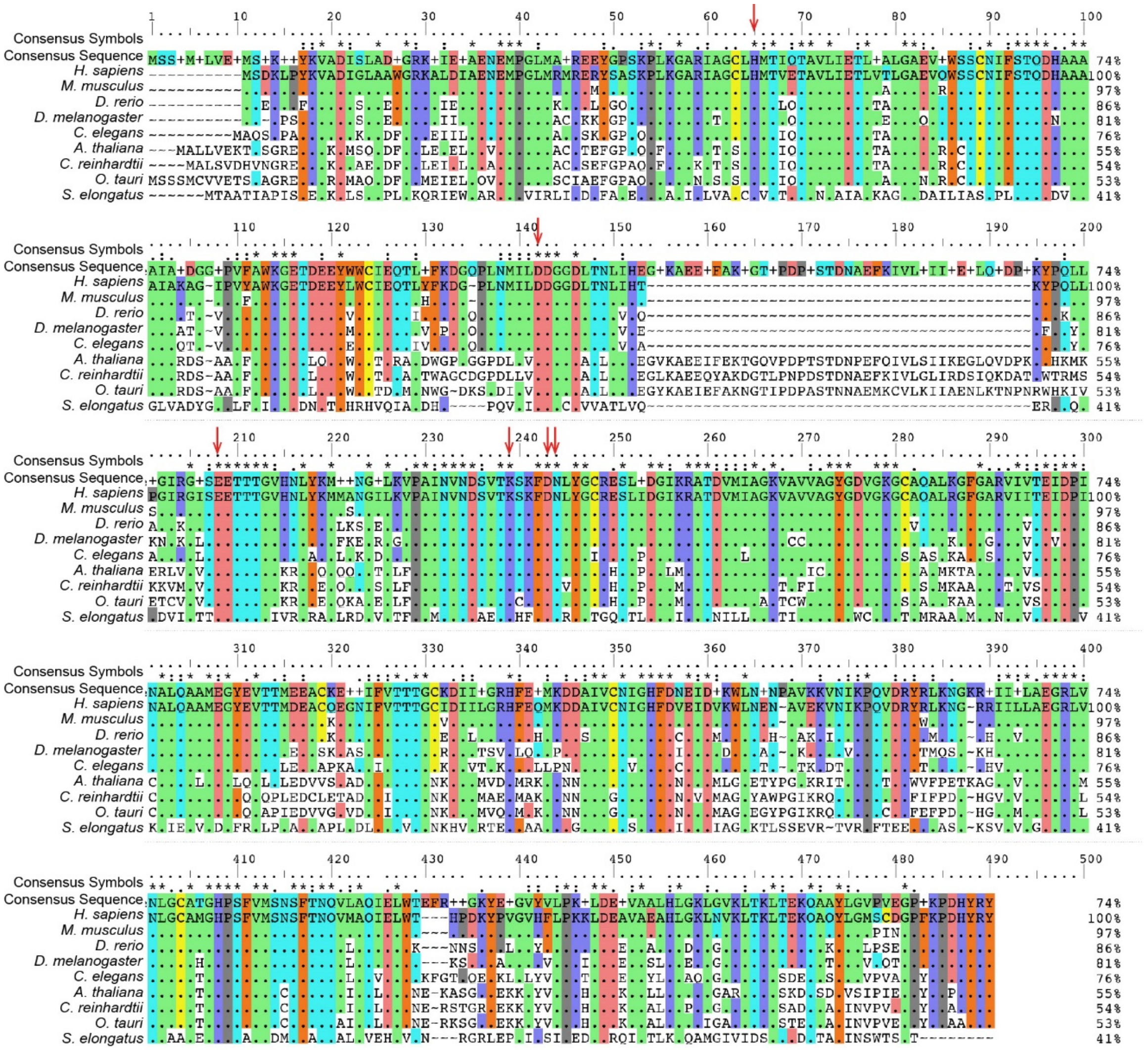


**Figure S1: Sequence alignment of full-length AHCY, related to Fig. 1.**
When amino acids are identical to human, a dot is shown in the alignment. Sequence identities (%), relative to human AHCY, are given on the right. The consensus symbols are shown on top (* = fully conserved residue, : = conservation of strongly similar properties, . = conservation of weakly similar properties). Red arrows indicate the residues that have been reported to be essential for rat AHCY function. The source of sequences is, for human, Genbank: NP_000678.1; mouse, Genbank: NP_057870.3; zebrafish, Genbank: NP_954688.1; fruit fly, Genbank: NP_511164.2; *C. elegans*, Genbank: NP_491955.1; *Arabidopsis*: NP_193130.1; *Chlamydomonas*, Genbank: XP_001693339.1; *Ostreococcus*, Genbank: XP_022839640.1; cyanobacteria, Genbank: WP_011243218.1.

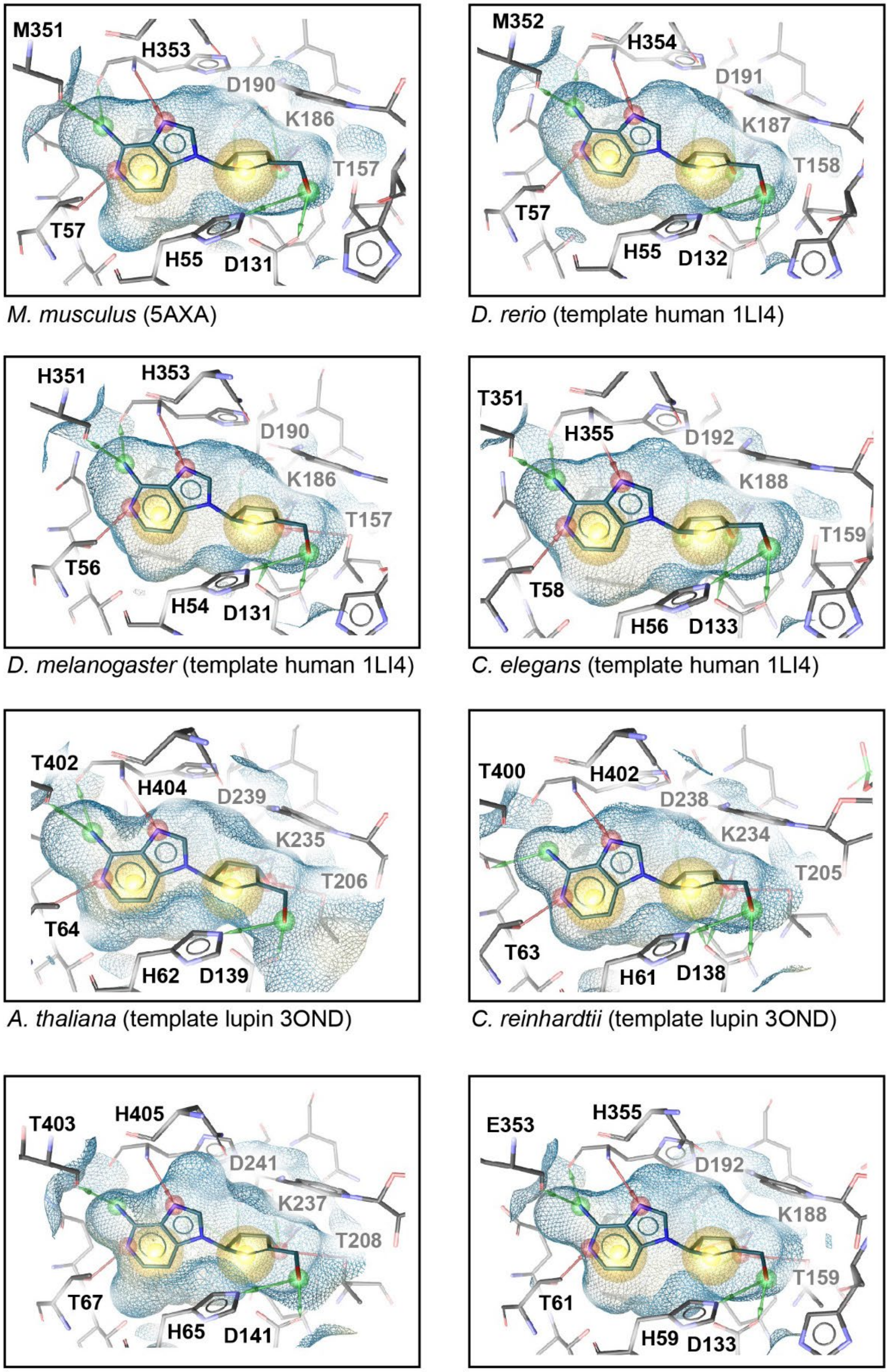


*M. musculus* (5AXA)

*D. rerio* (template human 1LI4)

*D. melanogaster* (template human 1LI4)

*C. elegans* (template human 1LI4)

*A. thaliana* (template lupin 3OND)

*C. reinhardtii* (template lupin 3OND)

*O. tauri* (template lupin 3OND)

*S. elongatus* (template human 1LI4)

**Figure S2: Molecular docking simulations of AHCY with DZnep, related to Fig. 1.**

Docking simulations of AHCY with DZnep, for each species as indicated below each picture, based on a published template crystal structure also indicated. The amino acids involved in DZnep binding are indicated together with their position. Note their conservation. The M351 in mouse (or human, see Fig. 1) is involved in DZnep binding via its backbone, explaining why, although this residue appears at first not conserved (H in fly, T in worm, plants and algae, and E in cyanobacteria), its conserved position in the active site is important. Red and green arrows are hydrogen bonds, yellow spheres are hydrophobic effects. Hydrogen atoms are not shown. The estimated free energies of binding for depicted DZnep docking conformations in kcal/mol were -9.92 for *M. musculus*, -9.50 for *D. rerio*, -9.29 for *D. melanogaster*, -9.46 for *C. elegans*, -9.50 for *A. thaliana*, -9.26 for *C. reinhardtii*, -9.68 for *O. tauri*, and -9.43 for *S. elongatus*.

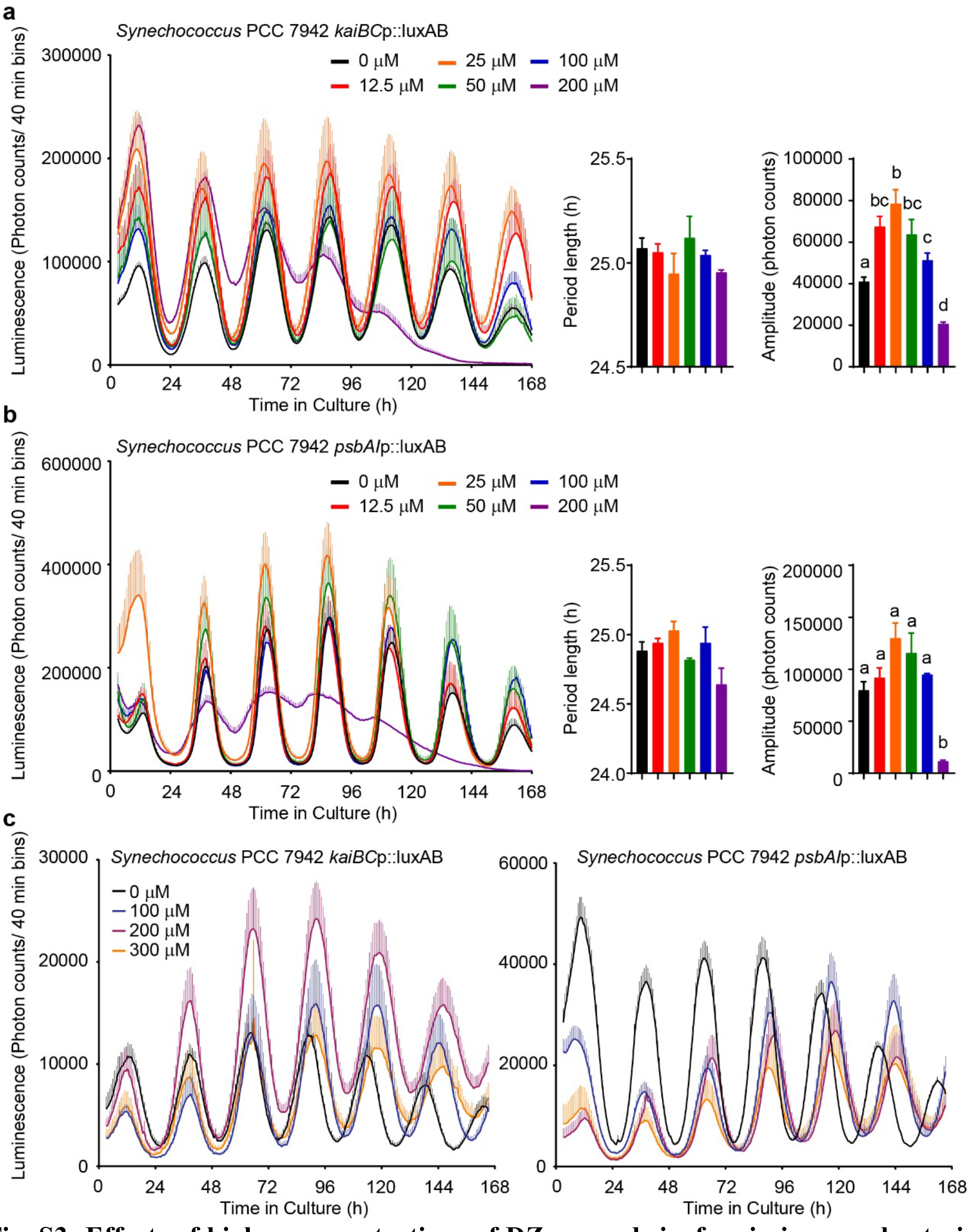


**Fig. S3: Effects of higher concentrations of DZnep and sinefungin in cyanobacteria, related to Fig. 5.**

**(a)** Left panel shows mean luminescence +/- SEM of *Synechococcus* PCC 7942 *kaiBC*p::luxAB knock-in strain, n = 3. Middle panel shows mean period +/- SEM of n = 3. Right panel shows mean amplitude +/- SEM of n = 3. **(b)** Same as (a) but using *Synechococcus* PCC 7942 *psbAI*p::luxAB knock-in strain. **(c)** Mean luminescence +/- SEM (n = 3) of *Synechococcus* strains treated with increasing concentrations of sinefungin. All bar graphs analyzed by One-Way ANOVA followed by Bonferroni post-hoc test, all comparisons at least $p < 0.05$.

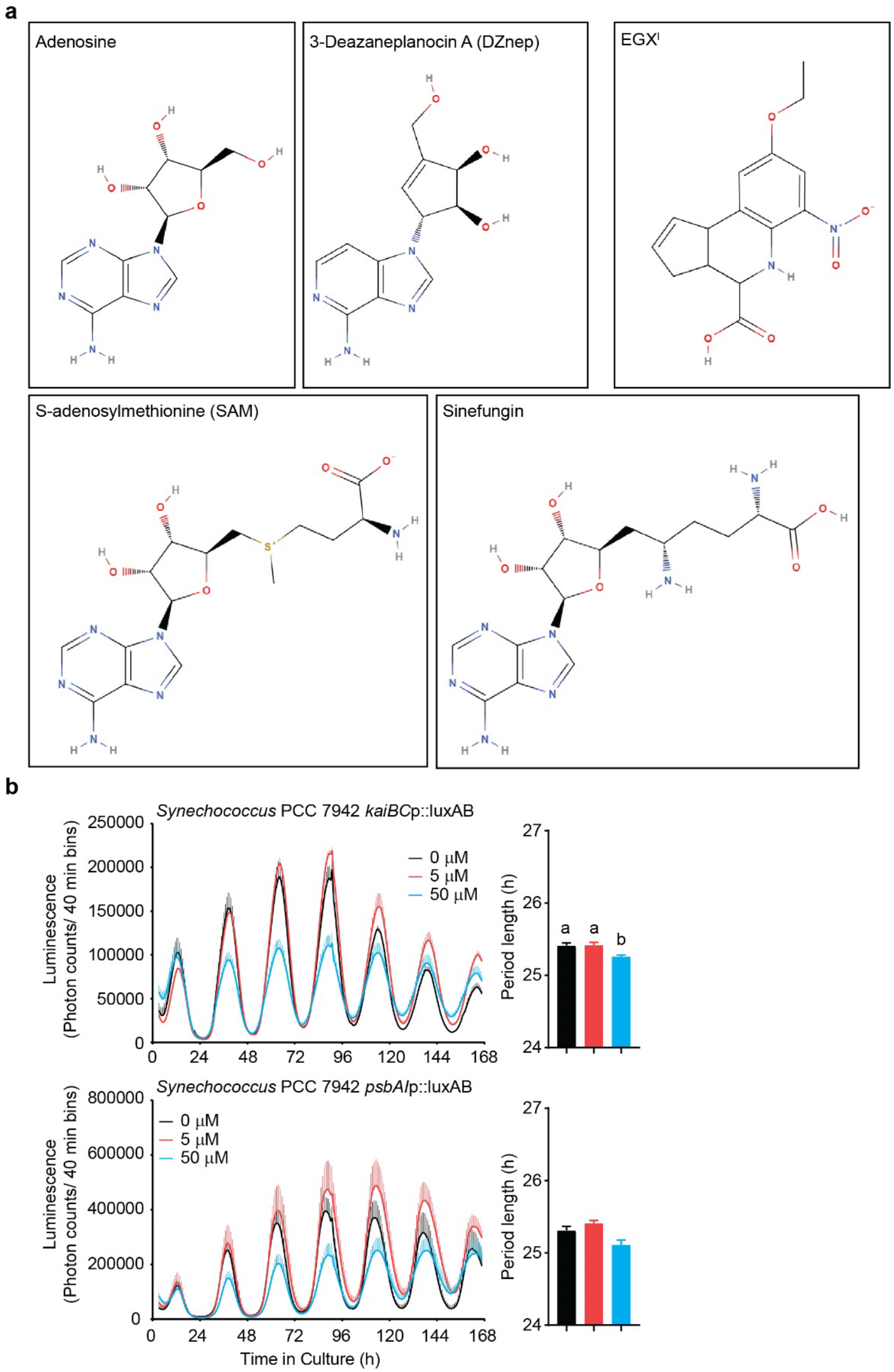


**Fig. S4: Comparison between the structure and effects of different methylation inhibitors used, related to Fig. 5.**

**(a)** Molecular structure of adenosine and its analogue DZnep, which inhibits methylation by binding to the adenosine binding-pocket of AHCY; SAM and its analogue sinefungin, which inhibits methylation by directly binding to methyltransferases; $EGX^I$, a cyclopentaquinoline carboxylic acid selective bacterial DNA methyltransferase inhibitor. **(b)** Upper left panel shows mean luminescence +/- SEM of Synechococcus PCC 7942 *kaiBC*p::luxAB knock-in strain treated with different concentrations of $EGX^I$, n = 3, with only the upper section of the error bars shown for clarity. Right panel shows mean period +/- SEM, n = 3. Lower panels show similar data but using *Synechococcus* PCC 7942 *psbAI*p::luxAB knock-in strain. All bar graphs analyzed by One-Way ANOVA followed by Bonferroni post-hoc test; all indicated comparisons at least $p < 0.05$.

**Captions for movies S1 to S3**

**Movie S1**: Animated structural superposition of AHCY from the 9 organisms investigated here, using human (1LI4), mouse (5AXA) or lupin (3OND) crystal structures as templates. The blue loop is specific to plants and green algae; DZnep is shown in yellow, $NAD^+$ in grey.

**Movie S2**: Time-lapse luminescence recordings of one representative embryo for each treatment.

**Movie S3**: Time-lapse luminescence recordings of one representative embryo for each treatment, with luminescence shown as a pseudo-color green and merged with brightfiled micrographs. The red arrows indicate the appearance of new somites.

**Supplementary References**


1. Yang X*, et al* (2003) Catalytic strategy of S-adenosyl-L-homocysteine hydrolase: Transition-state stabilization and the avoidance of abortive reactions. *Biochemistry* 42(7): 1900-1909.

2. Kusakabe Y*, et al* (2015) Structural insights into the reaction mechanism of S-adenosyl-L-homocysteine hydrolase*. Sci Rep* 5: 16641.

3. Sastry GM, Adzhigirey M, Day T, Annabhimoju R & Sherman W (2013) Protein and ligand preparation: Parameters, protocols, and influence on virtual screening enrichments. *J Comput Aided Mol Des* 27(3): 221-234.

4. Brzezinski K, Dauter Z & Jaskolski M (2012) High-resolution structures of complexes of plant S-adenosyl-L-homocysteine hydrolase (lupinus luteus)*. Acta Crystallogr D Biol Crystallogr* 68(Pt 3): 218-231.

5. Waterhouse A*, et al* (2018) SWISS-MODEL: Homology modelling of protein structures and complexes*. Nucleic Acids Res* 46(W1): W296-W303.

6. Benkert P, Biasini M & Schwede T (2011) Toward the estimation of the absolute quality of individual protein structure models*. Bioinformatics* 27(3): 343-350.

7. Van Der Spoel D*, et al* (2005) GROMACS: Fast, flexible, and free. *J Comput Chem* 26(16): 1701-1718.

8. Lindorff-Larsen K*, et al* (2010) Improved side-chain torsion potentials for the amber ff99SB protein force field. *Proteins* 78(8): 1950-1958.

9. Sousa da Silva AW & Vranken WF (2012) ACPYPE - AnteChamber PYthon parser interfacE. *BMC Res Notes* 5: 367-0500-5-367.

10. Morris GM*, et al* (2009) AutoDock4 and AutoDockTools4: Automated docking with selective receptor flexibility. *J Comput Chem* 30(16): 2785-2791.

11. Wolber G & Langer T (2005) LigandScout: 3-D pharmacophores derived from protein-bound ligands and their use as virtual screening filters. *J Chem Inf Model* 45(1): 160-169.

12. Wolber G, Dornhofer AA & Langer T (2006) Efficient overlay of small organic molecules using 3D pharmacophores. *J Comput Aided Mol Des* 20(12): 773-788.

13. Larkin MA*, et al* (2007) Clustal W and clustal X version 2.0. *Bioinformatics* 23(21): 2947-2948.

14. Lewis JA & Fleming JT (1995) Basic culture methods. *Methods Cell Biol* 48: 3-29.

15. Goya ME, Romanowski A, Caldart CS, Benard CY & Golombek DA (2016) Circadian rhythms identified in caenorhabditis elegans by in vivo long-term monitoring of a bioluminescent reporter. *Proc Natl Acad Sci U S A* 113(48): E7837-E7845.

16. Zielinski T, Moore AM, Troup E, Halliday KJ & Millar AJ (2014) Strengths and limitations of period estimation methods for circadian data. *PLoS One* 9(5): e96462.
